# Tryptanthrin Analogs Substoichiometrically Inhibit Seeded and Unseeded Tau4RD Aggregation

**DOI:** 10.1101/2024.02.02.578649

**Authors:** Ellie I. James, David W. Baggett, Edcon Chang, Joel Schachter, Thomas Nixey, Karoline Choi, Miklos Guttman, Abhinav Nath

## Abstract

Microtubule-associated protein tau is an intrinsically disordered protein (IDP) that forms characteristic fibrillar aggregates in several diseases, the most well-known of which is Alzheimer’s disease (AD). Despite keen interest in disrupting or inhibiting tau aggregation to treat AD and related dementias, there are currently no FDA-approved tau-targeting drugs. This is due, in part, to the fact that tau and other IDPs do not exhibit a single well-defined conformation but instead populate a fluctuating conformational ensemble that precludes finding a stable “druggable” pocket. Despite this challenge, we previously reported the discovery of two novel families of tau ligands, including a class of aggregation inhibitors, identified through a protocol that combines molecular dynamics, structural analysis, and machine learning. Here we extend our exploration of tau druggability with the identification of tryptanthrin and its analogs as potent, substoichiometric aggregation inhibitors, with the best compounds showing potencies in the low nanomolar range even at a ∼100-fold molar excess of tau4RD. Moreover, conservative changes in small molecule structure can have large impacts on inhibitory potency, demonstrating that similar structure-activity relationship (SAR) principles as used for traditional drug development also apply to tau and potentially to other IDPs.

## Introduction

Intrinsically disordered proteins (IDPs) play important structural and signaling roles *in vivo*.^1^ IDPs lack a single stable folded structure and often fulfill their roles by dynamically rearranging to interact with multiple different protein or macromolecular binding partners.^2,3^ This conformational plasticity also makes IDPs prone to misfolding and aggregation. Aggregation of specific IDPs into amyloid-type fibrils and a variety of smaller cytotoxic species is a hallmark of proteopathic diseases such as Alzheimer’s disease (AD), Parkinson’s disease, and Type II diabetes mellitus.^4–7^

Microtubule-associated protein tau, an IDP that stabilizes neuronal axons, exhibits pathological dysfunction and aggregation in a group of ∼20 neurodegenerative diseases that are collectively called tauopathies. Tau consists of six isoforms in the human brain that vary in their number of N-terminal inserts and repeat sequences in the microtubule binding region (MTBR). In primary tauopathies such as Pick’s disease, frontotemporal dementia with parkinsonism in chromosome 17, and progressive supranuclear palsy, amyloid deposition of one or more tau isoforms is the predominant pathological characteristic. Secondary tauopathies such as AD and Lewy body dementia (LBD) are characterized by the extensive deposition of both tau and one or more other aggregation-prone proteins (e.g., amyloid-β [Aβ] in AD, alpha-synuclein in LBD).^5,8–12^

Tauopathies and related neurodegenerative proteinopathies present a stark challenge to human health, affecting tens of millions globally, with very few effective therapeutic options.^8–13^ However, recently developed anti-Aβ therapeutic antibodies for AD can slow disease progression, albeit modestly and with a substantial risk of side effects.^14,15^ While these treatments are far from ideal, they do demonstrate that disease-modifying therapies targeting protein aggregation are indeed feasible, further supporting the idea that inhibiting tau aggregation could advance the treatment of both primary and secondary tauopathies. In particular, small molecule tau aggregation inhibitors could be made orders of magnitude more bioavailable in the brain than antibody drugs,^16^ and are better suited to target intracellular tau species. Accordingly, there is active interest in developing both antibody-based and small molecule tau aggregation inhibitors.

Despite this interest, tau and other IDP targets present unique challenges to rational drug design. First, the lack of well-defined structure in the physiological state prevents conventional structure-based discovery and design. Second, the specificity and strength of structure-activity relationships (SAR) for IDP/small molecule interactions remain open questions. For folded proteins, it is axiomatic that the chemical structure of a candidate therapeutic can be systematically modified to improve activity (e.g., selectivity, inhibition) by altering the physicochemical interaction between the ligand and protein target. It remains to be seen whether and how altering the 3D conformation of a small molecule modifies activity towards a disordered target.

Our approach differs from efforts to develop broad-spectrum aggregation inhibitors, whether based on natural products,^17,18^ molecular tweezer^19^ or designed peptide scaffolds,^20–22^ that target later oligomeric or fibrillar states. Instead, we target disordered, monomeric states with the goals of (1) achieving specificity for tau vs. other aggregation-prone proteins, and (2) interrupting aggregation before cytotoxic oligomers are populated.^23,24^ In previous work, we developed a novel enhanced sampling technique, repeated simulated annealing molecular dynamics (ReSA-MD), to generate a library of tau4RD conformers.^25^ A subset of these conformers displaying locally persistent structure was used as docking targets for *in silico* compound screening. To date, we have identified a novel family of tau4RD aggregation inhibitors, and a second family of compounds that potently binds to tau4RD fibrils without affecting the kinetics of aggregation.^25,26^ With this discovery pipeline, we now identify the tryptanthrin (TA) family as potent, substoichiometric inhibitors of tau4RD aggregation. Though TA, a plant alkaloid, is known to have anti-microbial,^27–30^ anti-inflammatory,^31,32^ anti-fungal,^33,34^ and anti-cancer activity,^35–38^ our work is the first demonstration of TA as an anti-amyloid agent. Here we describe the SAR of TA analogs against tau4RD aggregation and identify the mechanism by which they inhibit aggregation.

## Results

### Design, synthesis, and screening of TA analogs

We began by mining a 284,463-compound in-house library using a regression model trained on our previous *in silico*/*in vitro* screening effort^21^ to identify compounds likely to modulate tau aggregation. Characterizing heparin-induced tau4RD aggregation at a 1:2 protein:compound molar ratio using a thioflavin T (ThT) fluorescence assay revealed that TA (**1**) and three of its analogs (**2**–**4**) were among the most potent inhibitors (Fig. 1). ThT is widely used as a probe of protein aggregation, but is neither perfectly specific nor selective for amyloid aggregates. Therefore, orthogonal label-free assays were used to confirm that TA analogs are *bona fide* aggregation inhibitors, increasing the amount of tau4RD remaining in solution at the end of the reaction (Supplementary Figure S1). Several other chemically distinct screened compounds showed reasonable aggregation inhibition (compounds **S1**–**S20**; Supplementary Figure S2), further demonstrating the value of ReSA-MD-guided screening in targeting disordered proteins such as tau. However, this study focuses on the TA scaffold, both because these compounds displayed the greatest activity, and because they afforded an excellent opportunity to explore the structure-activity relationship of a novel family of tau aggregation inhibitors.

**Figure 1:**
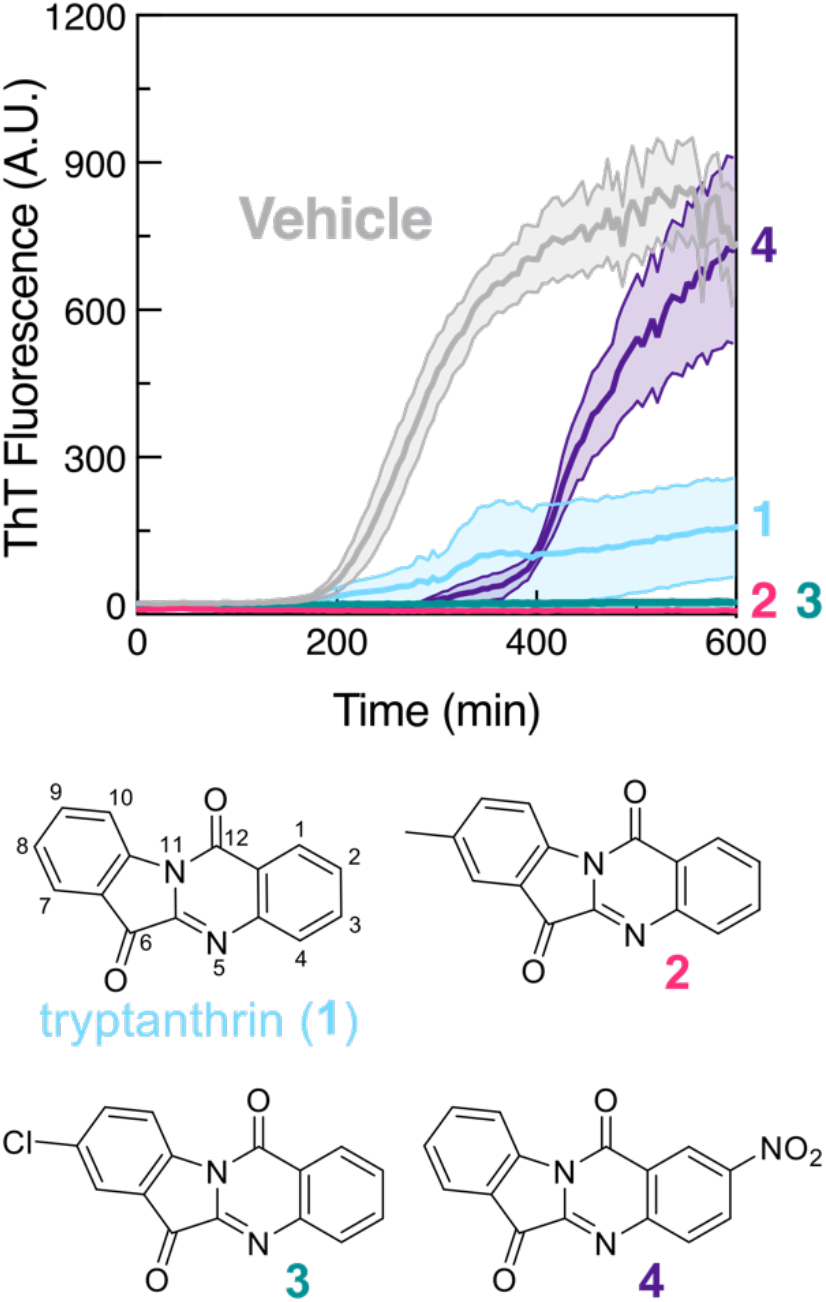
Inhibition of heparin-induced tau4RD aggregation by tryptanthrin and analogs, discovered by computationally-guided screening. Tau4RD (5 μM) aggregation in the presence of 5 μM 3 kDa heparin, with 5 μM of TA, analogs **2**–**4**, or vehicle control. (*n* = 3, error bands show mean +/-SEM.)

A 14-compound library (**5**–**18**) was synthesized based on 8-chloroTA (**3**), the most potent of the first four compounds, exploring the effects of modifications to the quinazoline ring (Fig. 2A). Synthesis from commercially available precursors used a one-step process driven by visible light in the presence of the dye Rose Bengal.^39^ The activity of **1** synthesized in-house was comparable to that of samples obtained from two different commercial vendors (Fig S3). All of these compounds strongly (10 of 13 compounds: **5, 6, 8, 10, 11, 13**–**18**) or moderately (3 of 13: **7, 9, 12**) inhibited tau4RD aggregation (Fig. 2B, C) at 5 μM. A subsequent 12 compounds (**19**–**30**) were synthesized based on 1,8-dichloroTA (**16**), selected based on its inhibitory properties and relatively low levels of cytotoxicity (Fig S4), to explore the effect of modifications to the indole ring. Three compounds (**20, 24, 25**) were dropped due to solubility issues, but seven of the remaining nine (**19, 22, 23, 27**–**30**) inhibited tau aggregation more strongly than **3**.

**Figure 2:**
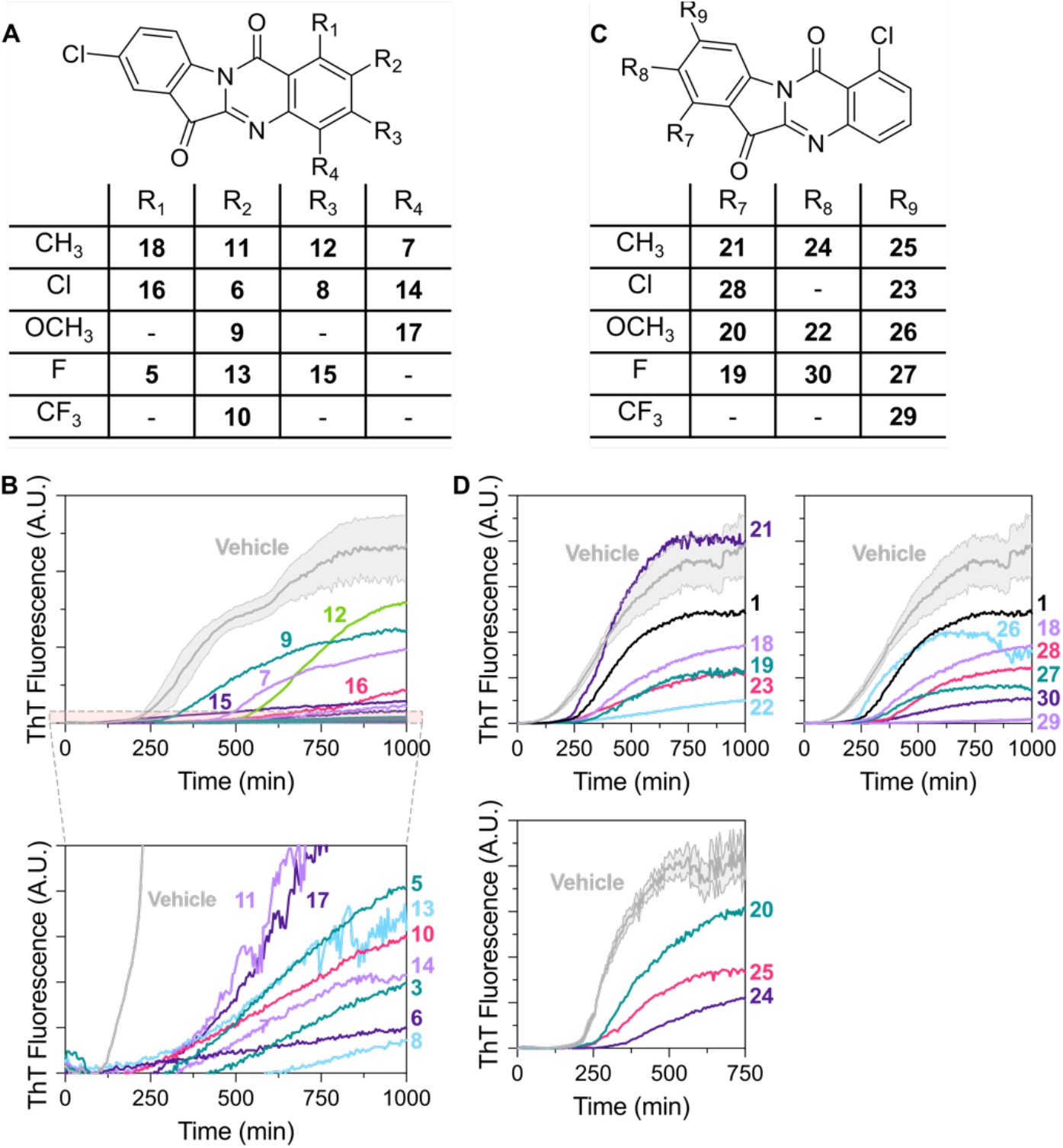
Effects of second and third generation TA analogs on heparin-induced tau4RD aggregation. A) Structures of second generation TA analogs **5**-**17**, with the table indicating substituents present at specified positions on the quinazoline ring for each analog. B) Tau4RD aggregation under standard conditions in the presence of second generation TA analog (5 μM) or vehicle control. The means (*n* = 3 or 4 technical replicates) are displayed for each trace. For visual clarity, error bands (mean +/-SEM) are displayed only for the vehicle trace. Other traces showed similar or lower technical variability. C) Structures of third generation analogs **19–30** derived from **16**. D) Tau4RD aggregation in the presence of third generation TA analog (5 μM) or vehicle control. Traces and error bands are formatted as in panel B. Compounds **20, 24**, and **25** (lower panel) were conducted at a DMSO concentration of 0.25% v/v, while all other experiments were conducted at 0.1% v/v DMSO.

### Potency of TA analogs

Evaluating and quantifying the potency of aggregation inhibitors from dose response experiments requires consideration of certain experimental design factors. The first is the ratio of inhibitor to other species involved in aggregation: in this case, tau4RD and the aggregation inducer heparin. Dose response aggregation assays yield a half-maximal effective concentration *EC*_50_, distinct from a dissociation constant *K*_*D*_ or an inhibitor binding constant *K*_*I*_. The *EC*_50_ would be expected to change if either the tau4RD or heparin concentrations were altered. It is important to recognize that the ratio of the *EC*_50_ to tau4RD concentration is more relevant in this context than the *K*_*D*_ or *K*_*I*_ itself. A stoichiometric inhibitor (i.e., one that must bind 1:1 to tau4RD for activity) would have a *EC*_50_ equal to the tau4RD concentration regardless of how low its *K*_*D*_ for tau4RD might be. Dose response assays were conducted at a concentration of 5 μM tau4RD and 5 μM heparin (with an average molecular weight of 3 kDa), in the presence of 1% DMSO, while compound concentrations varied from 62.5 nM to 5 μM.

A second factor is the choice of parameter used to quantify compound effects. TA analogs were seen to affect both the midpoint of the aggregation time course (*t*_50_) and the maximal fluorescence achieved at the end of the reaction (*F*_*plateau*_). Accordingly, we quantify dose response based on the ThT fluorescence at the *t*_50_ of the *vehicle* trace (*F*_50_; Fig. 3). This approach captures compound effects on both *t*_50_ and *F*_*plateau*_.

**Figure 3:**
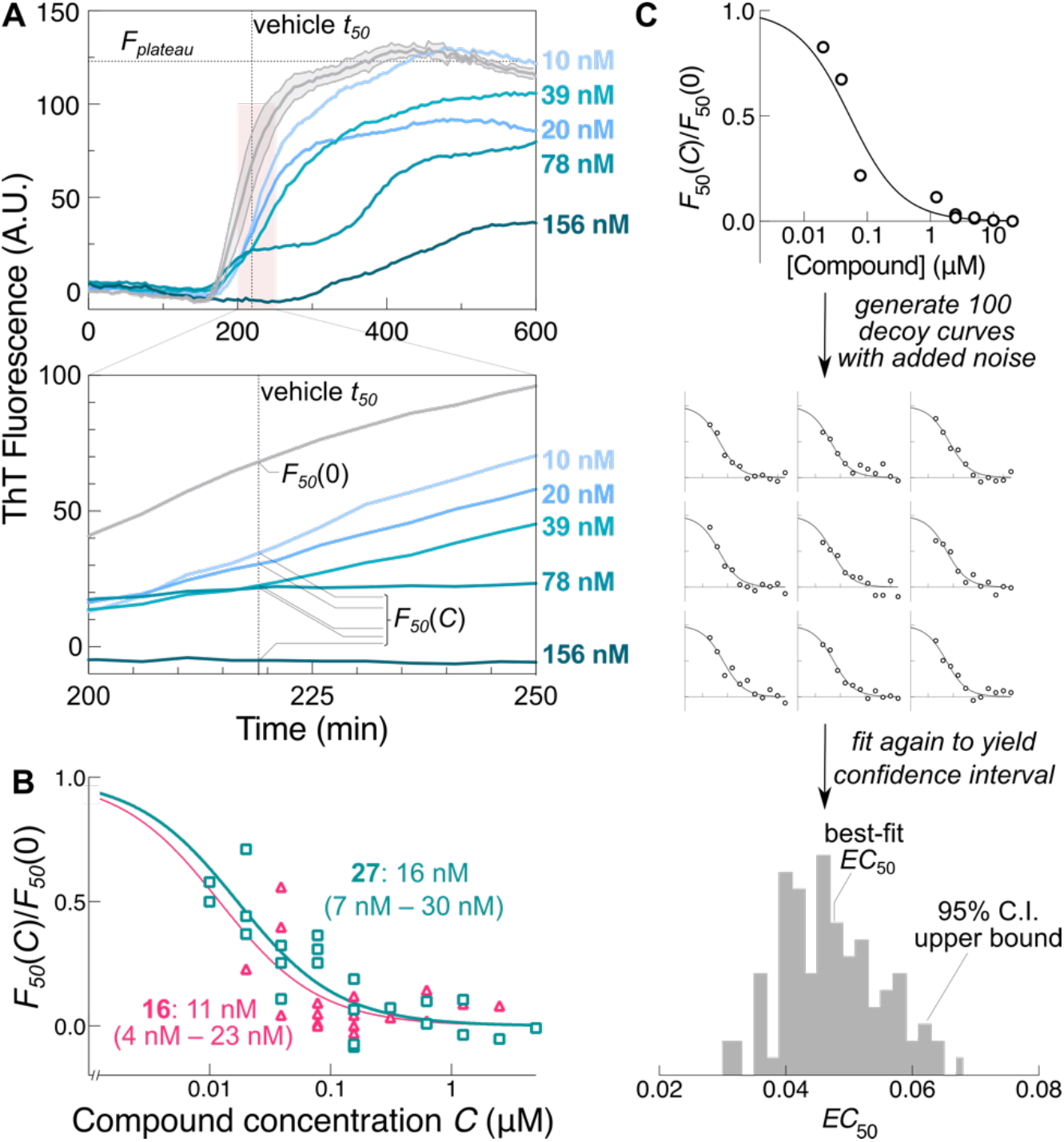
TA analogs inhibit tau4RD aggregation in a substoichiometric, dose-dependent manner. A) Tau4RD aggregation under standard conditions in the presence of varying concentrations of the third generation compound **27** or vehicle control (*n* = 3 or 4, mean displayed for all analog traces, S.E.M. error band displayed solely for the vehicle trace). *F*_50_ is defined as the fluorescence measured at the transition midpoint of the vehicle trace (*t*_50_), whether in the absence [*F*_50_(0)] or presence [*F*_50_(*C*)] of varying concentrations of compound. B) Dose-response curves of *F*_50_(*C*) normalized to *F*_50_(0) measured on the same plate for compounds **16** and **27**, demonstrating nanomolar potency. Best-fit *EC*_50_ values are shown, with 95% confidence intervals estimated from resampling shown in parentheses. C) Schematic of resampling protocol used to estimate confidence intervals of best-fit *EC*_50_ values.

A third consideration is accurately estimating the uncertainty in *EC*_50_ values obtained from dose response assays. Many sources of error can contribute to this uncertainty, and not all of them are normally distributed. Accordingly, a resampling procedure was used to more rigorously capture the range of parameter values consistent with observations. Briefly, a quadratic dose response equation (Eq. 2) was used to fit *F*_50_ vs. compound concentration data. Then, 100 synthetic ‘decoy’ dose response curves were generated using the best-fit values of *EC*_50_ with random noise added to match experimentally observed scatter in *F*_50_ values. The synthetic data sets were then re-fit using Eq. 2, and confidence intervals were then determined from the resulting distribution of re-fit *EC*_50_ values. We report the best-fit *EC*_50_ value from the experimental data, along with the 95% confidence interval, as a conservative estimate of the potency of each of the TA analogs. Equation 2 properly accounts for arbitrarily potent inhibition, and also yields as an additional global parameter *T*_*target*_, the concentration of species within the tau4RD population ensemble targeted by TA-family compounds.

Remarkably, dose-response studies revealed that many of these compounds are active at nanomolar concentrations even when tau is present in a significant molar excess. For example, **16** and **27** both have *EC*_50_ values below 100 nM in our assay (11 nM and 16 nM, with 95% C.I. upper bounds of 23 nM and 30 nM respectively). In other words, these compounds are able to block aggregation even when tau4RD is present in over a 100-fold molar excess. This clear substoichiometric activity suggests that TA analogs may be able to selectively target a sub-population of tau species that drive aggregation. *T*_*target*_ (2.3 nM, 95% C.I. upper bound 22 nM) is also well below the concentration of tau4RD, consistent with substoichiometric activity. The mechanistic implications of this selective engagement with particular protein species are explored below.

Furthermore, TA and its analogs exhibit strong structure-activity relationships. Conservative changes to the substituents of the quinazoline ring can either abrogate inhibitory activity (for example, compare **11** to positional isomers **7** and **12**) or enhance it (e.g., **16** vs. **6, 8** and **14**). Similarly, comparing **22** vs. **26** or **29** vs. **23** illustrates the effects that positional isomers and substitutions on the indole ring can have on activity. This pattern of SAR indicates that the interactions between TA analogs and tau4RD are specific and rationalizable, despite the intrinsic disorder of tau4RD, which is consistent with our previous work.^22^

### Mechanism of aggregation inhibition by TA analogs

Given the potency and substoichiometric inhibition of tau4RD aggregation demonstrated by TA-family compounds, we turned our attention to understanding the mechanism driving their activity. Our design and discovery strategy targets monomeric states of tau4RD, meaning that TA analogs would be expected to target the earliest stages of the aggregation pathway. To test this idea, we performed delayed addition experiments. Briefly, instead of being included at the start of an aggregation reaction, TA analogs or vehicle were introduced at defined timepoints within the lag, elongation, and plateau phases of aggregation (Fig. 4). The delayed addition results show that TA analogs inhibited aggregation when added during the lag phase (where primary nucleation is the dominant process), but did not inhibit aggregation when added later in the elongation phase (where fibril growth and secondary nucleation predominate). From this data, we concluded that because TA analogs are more potent inhibitors when added immediately after aggregation initiation, TA analogs primarily act on tau4RD nucleation. The ability to selectively target nucleating species, or their precursors, would nicely explain the strongly substoichiometric activity of TA analogs.

**Figure 4:**
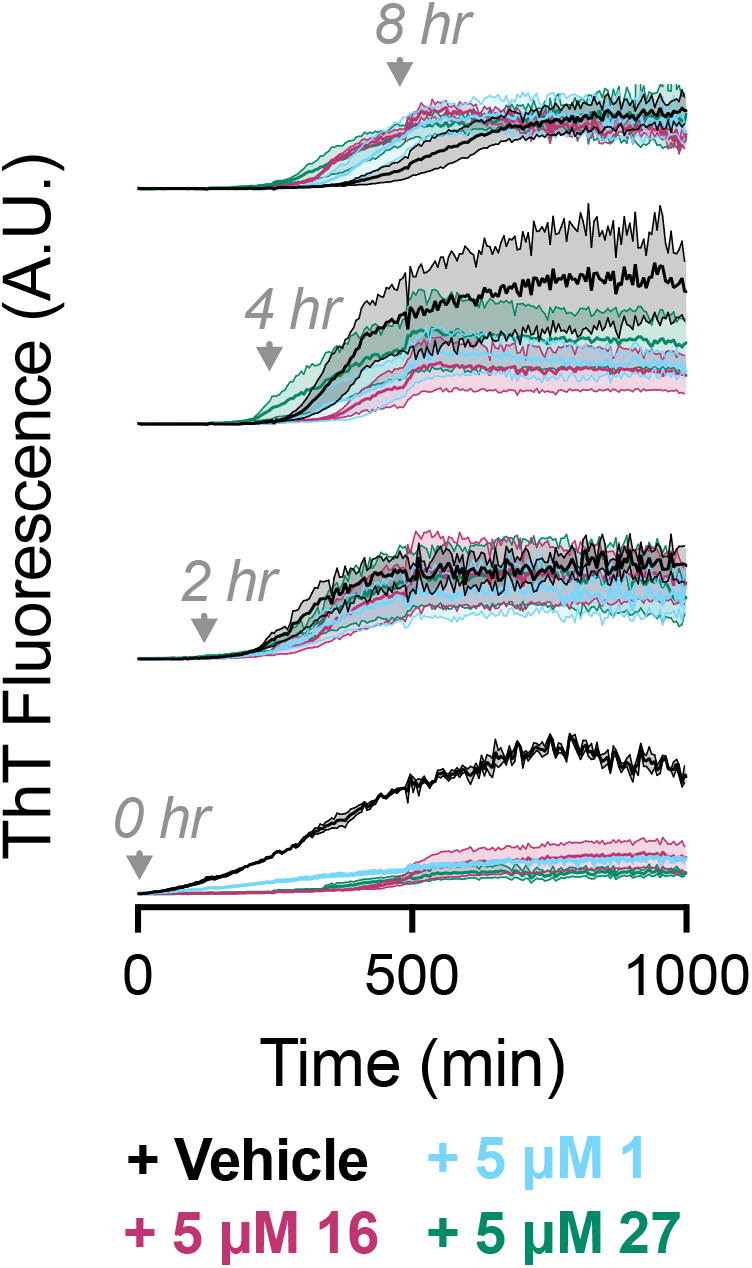
TA analogs are most active early in the aggregation process. Tau4RD aggregation with delayed addition of 5 μM TA analog at t = 0, 2, 4, or 8 hours (*n* = 3, mean +/-SEM). Inhibition was observed when analogs were added early in the aggregation lag phase but not once the elongation phase began. This suggests that TA analogs preferentially inhibit primary nucleation.

To directly determine whether TA analogs have any effect on the tau4RD elongation rate, we performed seeded aggregation assays. Seeding tau4RD with preformed fibrils bypasses the nucleation phase, allowing direct observation of elongation kinetics. We performed seeded aggregation assays with tau4RD:TA stoichiometries between 1:2 and 1:0.125. We observed a dose-dependent decrease in tau4RD elongation rate (*k*_*obs*_, AU/min) with increased stoichiometric equivalents of TA analogs (Fig. 5). The dose response of the observed elongation rate indicated that TA analogs inhibit seeded tau4RD aggregation with mid-nM potency.

**Figure 5:**
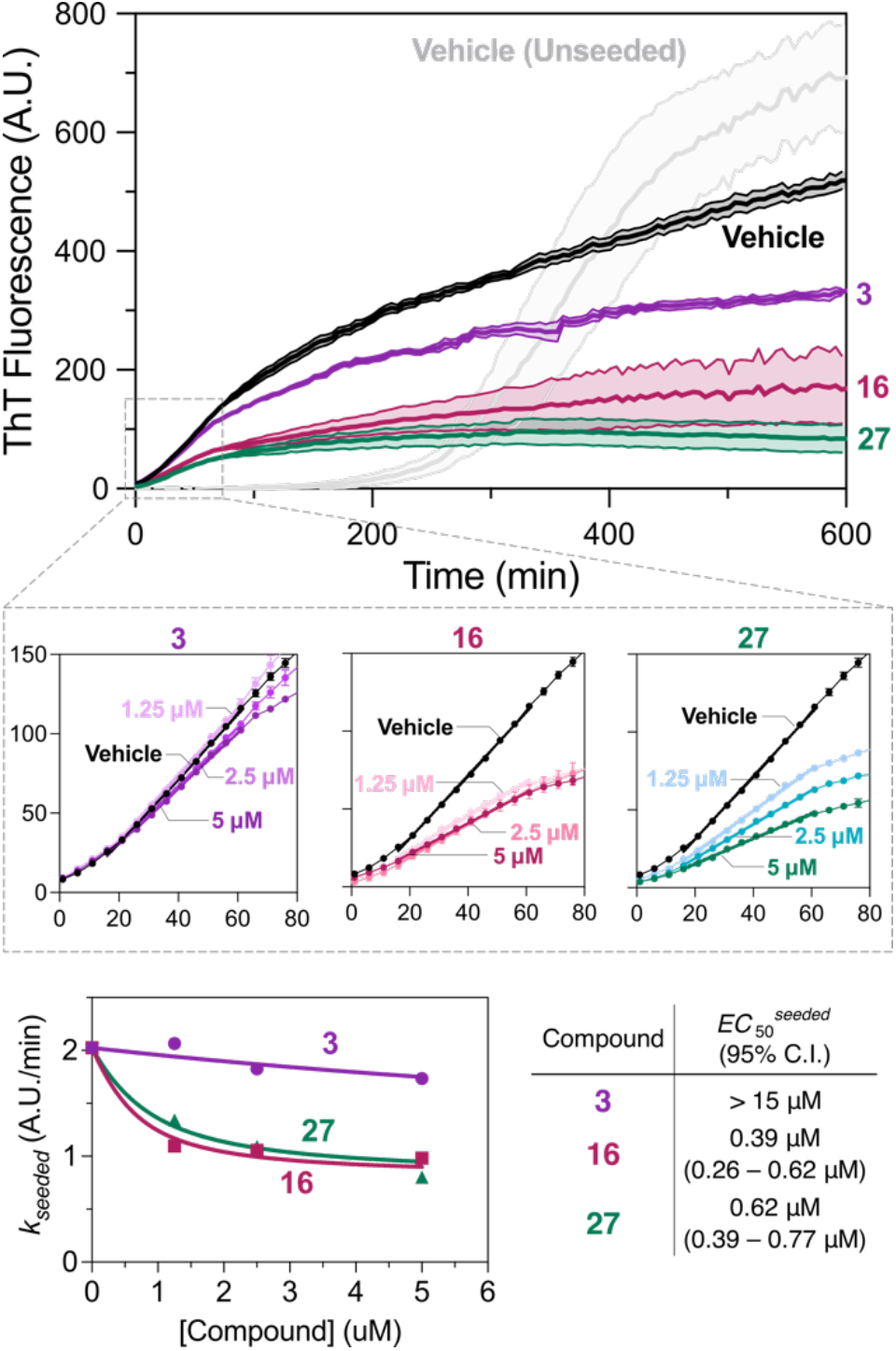
Inhibition of seeded tau4RD aggregation by TA analogs. A) Aggregation of 5 μM tau4RD and 5 μM 3 kDa heparin, seeded with 5% (w/w) pre-formed fibril in the presence of 5 μM TA analog or vehicle control. Traces indicate mean +/-SEM (*n* = 3 or 4). B) Seeded elongation measured over a range of compound concentrations, with linear fits (bold lines) used to estimate elongation rates. Traces indicate mean +/-SEM (*n* = 3 or 4). C) *k*_*seeded*_ rates from B plotted against compound concentration and fit to Equation 2, with 95% C.I. estimated by resampling.

**Figure 6:**
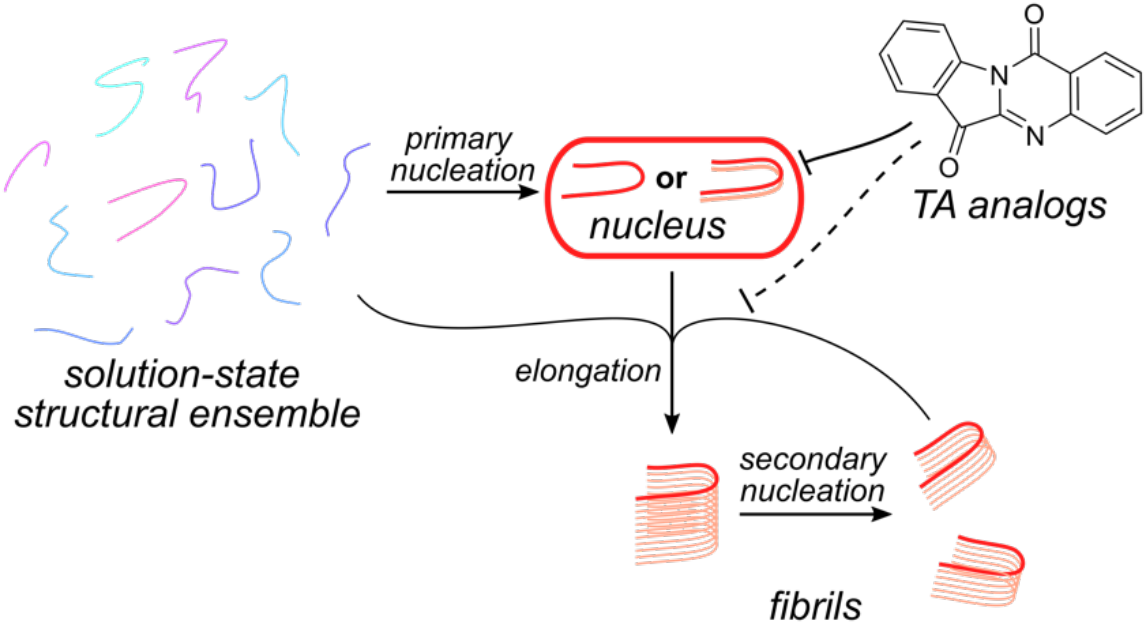
Proposed mechanism of action of tryptanthrin and its analogs. Substoichiometric inhibition of early-stage aggregation indicates that TA analogs bind selectively and tightly to tau aggregation nuclei or their close precursors. Substoichiometric inhibition of seeded aggregation indicates that TA analogs can also inhibit subsesquent elongation or secondary nucleation, albeit with lower potency.

## Discussion

The development of potent, specific small-molecule aggregation inhibitors for tau and other aggregation-prone IDPs remains a major open challenge. Our work here demonstrates that computationally-guided screening followed by focused analog design can, at least in this case, provide a solution. TA and its analogs are an attractive scaffold for further development for several reasons.

TA-family compounds display remarkable potency and substoichiometric activity, at levels that have never (as far as we are aware) been reported for small-molecule tau aggregation inhibitors. At least two compounds (**16** and **27**) are active at concentrations well below 100 nM, and inhibit unseeded aggregation of tau4RD at >50-fold molar excess. As discussed above, when it comes to aggregation inhibition, the substoichiometric nature of this activity is even more exciting than raw potency or binding affinity. For a target like tau, with a total concentration of ∼2 μM in the neuron,^8^ it would be much more tractable to deliver nM concentrations of a substoichiometric therapeutic to the brain than μM concentrations of a stoichiometric one.^40^

Moreover, TA-family compounds demonstrate that dramatic changes in activity can result from conservative changes to small-molecule structure, even with a target as disordered as tau. Firstly, this finding confirms that the interactions between tau and the small molecules presented here are specific in nature. Secondly, it suggests that iterative analog optimization is a feasible strategy to generate high-affinity ligands of disordered proteins, following similar paradigms well-established for structured protein targets.^41^

Mechanistic studies shed light on how exactly TA and its analogs modulate tau aggregation. TA-family compounds more potently inhibit unseeded than seeded tau4RD aggregation, with *EC*_50_ values for the latter higher by about an order of magnitude. (Note that the inhibition of seeded aggregation is still substoichiometric, with *EC*_50_s about 10-fold lower than tau4RD concentration.) In nucleation-dependent polymerization (NDP) models, seeded aggregation is driven by elongation (i.e., monomers adding on to existing fibrils) and secondary nucleation processes (by which existing fibrils generate new fibril growth sites).^42^ Unseeded aggregation is driven by primary nucleation (*de novo* formation of nuclei or fibrillar seeds from soluble material) in addition to elongation and secondary nucleation. Accordingly, the difference in potency towards unseeded and seeded reactions indicates that TA-family compounds preferentially target early stages of tau4RD aggregation. This is clearly backed up by delayed addition experiments that demonstrate a loss of activity if aggregation is allowed to continue unchecked for defined periods of time before compounds are added. Moreover, the strongly substoichiometric activity of TA analogs indicates that they must bind selectively and tightly to either the aggregation nucleus or a close precursor (whether monomeric or oligomeric). This raises the exciting prospect that characterizing TA analogs’ effects on tau conformation could shed light on these enigmatic species.

The discovery and functional characterization of novel TA inhibitors of tau aggregation positions us to pursue important future questions. On the one hand, it will be important to determine where TA analogs bind to tau and how they affect its conformation. Given the disordered nature of the target, X-ray crystallography or cryo-EM are unlikely to be useful,^2,3^ and multidimensional NMR would require intractably high concentrations of TA analogs in solution.

Accordingly, we are developing hydrogen/deuterium exchange mass spectrometry (HDX-MS) methods to characterize tau/small-molecule complexes.^43,44^ On the other hand, establishing the biological activity of TA-family compounds is the essential next step in their development as potential tau-targeting therapeutics. Studying their effects on tau aggregation in cells or *in vivo*, and their ability to slow the propagation of tau pathology from neuron to neuron, would reveal whether and how the intriguing *in vitro* activity of tryptanthrin and its analogs translates to the biological environment.

## Materials and Methods

### Protein Expression and Purification

Tau4RD was expressed and purified from *E. coli* BL-21 (DE3) as previously reported.^21^ Briefly, a plasmid containing the His-tagged tau4RD gene with a TEV cleavage site was gifted from the Rhoades lab at the University of Pennsylvania (Philadelphia, PA). To express protein, BL-21 cells containing the tau4RD gene were grown at 37 °C until the OD reached 0.6-0.7. The temperature was then lowered to 16 °C and protein expression was induced with the addition of Isopropyl B-D-1-thiogalactopyranoside to a final concentration of 0.4 mM. The culture was allowed to grow overnight, then cells were harvested by centrifugation. Expressions were resuspended, flash frozen, and stored at -80 °C in lysis buffer [50 mM Tris, 500 mM NaCl, and 10 mM imidazole (pH 8 at 4 °C)]. Upon thawing for purification, Halt Protease inhibitor cocktail (Life Technologies), DNase, RNase, and phenylmethanesulfonylfluoride in ethanol were added to cells before lysis with a French press. The lysate was centrifuged at 2800*g* for 45 minutes, then the supernatant was passed through 0.4 μm filters and loaded onto a 5 mL Ni-NTA agarose column. The protein was eluted by increasing the concentration of imidazole, then the eluent was dialyzed against a 10 μM imidazole buffer during an overnight incubation with 1 mM dithiothreitol and 100 μL of 8.64 mg/mL TEV protease (expressed in *E. coli*) at 4 °C. This solution was loaded onto a Ni-NTA column and the flow-through was collected and concentrated to 1-2 mL with a 3 kDa molecular weight cutoff centrifugal filter. The concentrate was fractionated using a 25 mL S200 extended gel-filtration column as previously described.^25^ Protein purity was confirmed by precast 12% Bis-Tris SDS-PAGE gels (Invitrogen). Protein was concentrated, aliquoted, and flash-frozen before storage at –80 °C. Protein concentrations were determined using the Pierce BCA Protein Assay Kit (Thermo Fisher Scientific).

### Aggregation Assays

#### Unseeded

All aggregation assays were performed at 37 °C and pH 7.4 in filtered (0.22 μm) aggregation buffer containing 20 mM Tris, 50 mM NaCl, 1 mM TCEP, 1 mM EDTA. EDTA was added to aggregation assays because the presence of trace divalent metal cations was found to affect tau4RD aggregation.^45,46^ Aggregation buffer was made fresh and passed through a 0.22 μm syringe filter before use. (Tau4RD, TA analogs, and heparin were not filtered because filtration of these components resulted in spurious aggregation results.) Aggregation assays were performed at 5 μM tau4RD, 5 μM ∼3 kDa heparin, and 5 μM TA analog unless otherwise indicated. DMSO concentration in all assays was 0.1% unless otherwise stated. Solutions of tau4RD and TA analogs (or an equivalent volume of DMSO) were prepared at 2.105x the desired final concentrations in aggregation buffer, and 95 μL aliquots of each were then added to the wells of a 96-well black flat-bottom non-binding plate (Corning). A 1 mM stock of ThT in aggregation buffer was passed through a 0.22 μm syringe filter, and then 10 μL of this solution was added to the wells for a final concentration of 50 μM ThT. Immediately before beginning fluorescent readings, 95 μL of 5 μM unfractionated ∼3 kDa heparin (MP Biomedicals, Irvine, CA) dissolved in aggregation buffer was added to each well, bringing the total well volume to 200 μL. The plate was covered with a piece of polyolefin or polypropylene sealing tape (Thermo Fisher Scientific) and monitored with a BioTek Synergy HTX plate reader (Agilent, Santa Clara, CA). Intrinsic fluorescence from TA analogs did not interfere with monitoring reaction progress by the increase in ThT fluorescence (λ_ex_ = 440 nm, λ_em_ = 485 nm). Reads were taken every 5 min after linear agitation (300 rpm) for 1 min. Experiments were performed with at least three technical replicates. Baseline correction was performed by subtracting the fluorescence value (A.U.) of the buffer/DMSO trace from experimental traces.

#### Seeded

Pre-formed tau4RD seeds were generated by harvesting tau4RD fibrils after 17 h of aggregation. Briefly, wells containing tau4RD and DMSO from unseeded aggregation assays were transferred to individual snap-cap tubes and centrifuged at 21,000*g* for 30 min at 4 °C. 100 μL of supernatant was discarded, then 100 μL of fresh aggregation buffer was added to each tube. Each 200 μL aliquot was sonicated on ice for 5 rounds of 2 sec (Cole-Parmer 4710 Series Ultrasonic Homogenizer, 50% output) to disrupt tau4RD amyloid fibrils. Seeds were added to aggregation assays at 5% (v/v). Aggregation assays were then performed as above, except that 10 μL of tau4RD seeds or aggregation buffer was added to the wells after addition of 85 μL of heparin.

#### Delayed addition

Delayed addition aggregation assays were prepared similarly to unseeded aggregation reactions except that DMSO and TA analogs were not included in the tau4RD aliquots introduced to the wells. TA analogs were diluted in DMSO to 1 mM and held until the appropriate time point. Tau4RD, ThT, and heparin were prepared as above and plated in the stated order. At t = 0, 120, 240, or 480 min, 1 μL of 1 mM TA analog or 1 μL of DMSO were pipetted into the appropriate wells to create three technical replicates of each condition. To do so, the plate was removed from the plate reader and unsealed at each time point, then resealed and replaced after the addition and allowed to continue incubating. Well volume was 201 μL after addition. Reaction progress was monitored as above.

### Gel Densitometry

Aggregation assays were stopped at the indicated time and wells were pipetted into individual snap-cap tubes. The tubes were centrifuged 21,000*g* for 30 min at 4 °C, after which 15 μL of supernatant was added to 3 μL of 5x SDS loading dye. Concentration standards were prepared at 5 μM, 3 μM, and 1 μM and t = 0 samples were prepared with 5 μM final concentration of both tau4RD and the respective TA analog. Samples were boiled for 45 sec, cooled on ice, and 15 uL was added to precast 12% Bis-Tris SDS-PAGE gels. Gels were imaged with a Li-Cor Odyssey CLx gel scanner and band intensity was determined using ImageJ.^47^ Experiments consisted of 3 technical replicates and the results were averaged (mean).

### EC_50_ Determination and Resampling

Unseeded aggregation assays were performed as above with 5 μM tau4RD and TA analog at concentrations between 62.5 nM and 10 μM. DMSO content was 0.25% for experiments involving **20, 24**, and **25** and 0.1% for experiments involving all other compounds. ThT fluorescence traces were baseline corrected and averaged. Non-linear least squares fitting in Prism 9.0 (GraphPad Software, Boston, MA) was used to determine *t*_50_, the midpoint of the aggregation curve, from the vehicle control trace from the following equation:

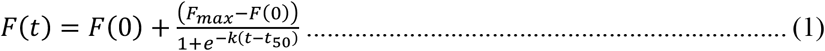

Here, *F*(0) is the baseline fluorescence at the start of the reaction, *F*_*max*_ is the maximal fluorescence at the plateau after aggregation is complete, and *k* is a rate parameter that describes the steepness of the transition.

We then define *F*_50_(*C*) as the ThT fluorescence in the presence of a given concentration *C* of a TA analog at the *t*_50_ (midpoint) *of the vehicle control trace. F*_50_ values measured over a range of compound concentrations were fit to the following quadratic dose-response equation to determine analog *EC*_50_:

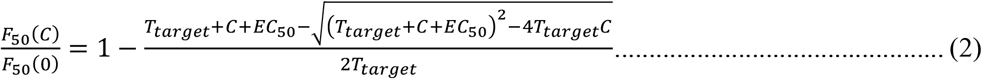

Here, *F*_50_(0) is the fluorescence of the vehicle control trace at its *t*_50_ (i.e., in the absence of compound), and *T*_*target*_ is the concentration of species within the tau4RD ensemble that are targeted by the compound. The value of *T*_*target*_ was globally shared across all dose response fitted curves.

To estimate the 95% confidence interval of the mean *EC*_50_ values, 100 synthetic “decoy” datasets were generated based on the best-fit values obtained from Equation 2. Gaussian noise equal to the RMSE observed in the *EC*_50_ fit was applied to the decoys which were each then fit to Equation 2. The upper and lower 95% confidence intervals were determined by binning the *EC*_50_ values of the decoy datasets.

### TA Synthesis, Purification, and Storage

Following the protocol of Hou et al., TA analogs were synthesized by reacting isatin derivatives (1.0 eq) with benzoxazine-2,4-dione derivatives (1.2 eq) in the presence of Rose Bengal (0.050 eq) and potassium carbonate (1.0 eq) at ambient temperature under illumination for 18h. TA analogs were purified by reverse-phase HPLC followed by supercritical fluid chromatography, characterized by ^1^H NMR and/or mass spectrometry, and lyophilized. Details on the synthesis, purification, and characterization of each TA analog are presented in Supplementary Material. DMSO stocks of TA analogs were prepared, aliquoted, and stored at –20 °C until use.

## Acknowledgments

This work was supported by the NIH (T32GM007750 to E.I.J.) and the Seattle Partnership for Research on Innovative Therapies (to A.N.). The authors gratefully acknowledge Elena Arroyo Holland and Mia Cervantes for assistance with protein expression, as well as Dr. Cathy Tralau-Stewart and Dr. C. Alex Goddard (Takeda Pharmaceuticals) for administrative support. The authors also gratefully acknowledge helpful discussions with Prof. Brian Kraemer, Prof. Tuomas Knowles, and Prof. Michele Vendrusculo.

## Author contributions

E.I.J., D.W.B. and A.N. conceived the project. E.I.J. performed *in vitro* aggregation assays. D.W.B., E.I.J. and A.N. performed the computational work. E.D.C., T.N. and K.C. were responsible for the synthesis and validation of TA analogs. J.S. was responsible for cytotoxicity assays. E.I.J., and A.N. drafted the manuscript, and all authors participated in editing.

