## Supplementary Material for "Tryptanthrin Analogs Substoichiometrically Inhibit Seeded and Unseeded Tau4RD Aggregation"

**Synthesis and Characterization of Tryptanthrin Analogs**

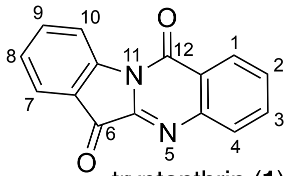
tryptanthrin (**1**)

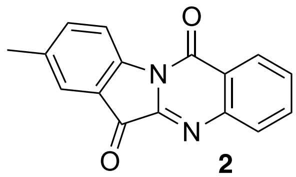
**2**

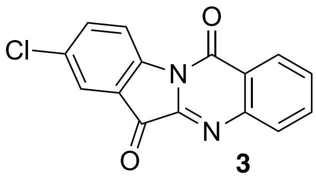
**3**

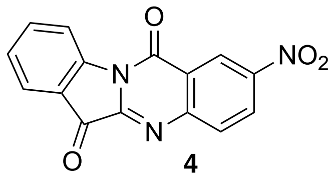
**4**

Compounds **1–4** were sourced from Axcelead Drug Discovery Partners (Fujisawa, Japan). Additional **1** was purchased from Sigma Aldrich (St. Louis, MO) and MedChemExpress (Monmouth Junction, NJ).

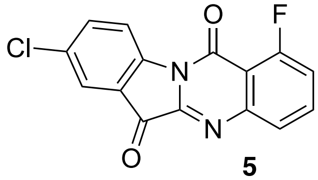
**5**

Compound **5** (8-chloro-1-fluoro-indolo[2,1-b]quinazoline-6,12-dione) was synthesized as follows: a solution of 5-fluoro-1H-3,1-benzoxazine-2,4-dione (1.2 eq, 26 mg, 0.13 mmol), Rose Bengal (0.050 eq, 5.6 mg, 0.0055 mmol) and DMA (0.9 mL) was added to a 4 mL vial containing 5-chloroisatin (1.0 eq, 20 mg, 0.11 mmol) and potassium carbonate (1.0 eq, 15 mg, 0.11 mmol). The resulting mixture was stirred (uncapped) at ambient temperature under a 100 lumen white LED light (Milwaukee Tool, Brookfield, WI) for 18h. DMSO (1 mL) was added to solubilize the mixture, which was then filtered and purified by HPLC (Phenomenex Gemini® C18, 5 µm, ID 30 mm x 150 mm, eluting with 10-90% acetonitrile (0.035% TFA)/water (0.05% TFA), and then dried *in vacuo*. The resulting solid was repurified by supercritical fluid chromatography (2-PIC column (30 x 150mm, 5 micron), eluting with 5-50% MeOH/$CO_2$:methanol, MeOH modified with 0.1% ammonium hydroxide) to give 8-chloro-1-fluoro-indolo[2,1-b]quinazoline-6,12-dione (2.1 mg, 6.4 % yield). ESI-MS m/z [M+H]+ calc'd for $C_{15}H_6ClFN_2O_2$ 300.0; found 300.9.

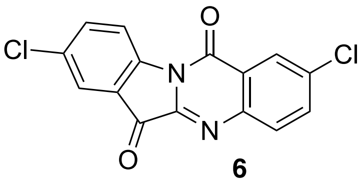
**6**

Compound **6** (2,8-dichloroindolo[2,1-b]quinazoline-6,12-dione) was synthesized (1.7 mg, 4.9 % yield) using 6-chloro-1H-3,1-benzoxazine-2,4-dione as a starting material and the analogous procedure described for Compound **5** above. $^{1}$H NMR (400 MHz, chloroform-*d*) δ ppm 7.76 - 7.80 (m, 1 H) 7.81 - 7.85 (m, 1 H) 7.89 - 7.92 (m, 1 H) 7.98 - 8.02 (m, 1H) 8.41 - 8.44 (m, 1 H) 8.59 - 8.63 (m, 1 H). ESI-MS $[M+H]^+$ calc'd for $C_{15}H_6Cl_2N_2O_2$, 316.0; found, 316.9.

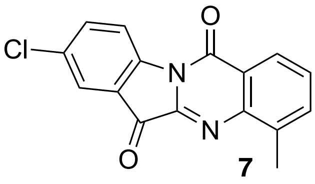

**7**

Compound 7 (8-chloro-4-methyl-indolo[2,1-b]quinazoline-6,12-dione) was synthesized (13 mg, 38 % yield) using 8-methyl-1H-3,1-benzoxazine-2,4-dione as a starting material and the analogous procedure described for Compound **5**. ESI-MS m/z [M+H]+ calc'd for $C_{16}H_9ClN_2O_2$ 296.0; found 297.0.

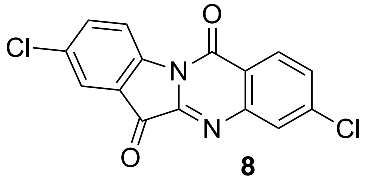

**8**

Compound **8** (3,8-dichloroindolo[2,1-b]quinazoline-6,12-dione) was synthesized (2.8 mg, 7.3 % yield) using 7-chloro-1H-3,1-benzoxazine-2,4-dione as a starting material and the analogous procedure described for Compound **5**. $^1$H NMR (400 MHz, chloroform-*d*) δ ppm. 7.65 - 7.69 (m, 1 H) 7.76 - 7.80 (m, 1 H) 7.89 - 7.92 (m, 1 H) 8.02 - 8.05 (m, 1 H) 8.37 - 8.41 (m, 1 H) 8.58 - 8.62 (m, 1 H) ESI-MS m/z [M+H]+ calc'd for $C_{15}H_6Cl_2N_2O_2$ 316.0; found 316.9.

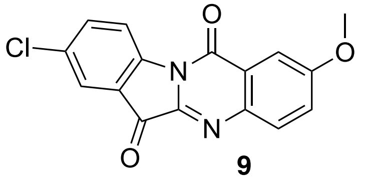

**9**

Compound **9** (8-chloro-2-methoxy-indolo[2,1-b]quinazoline-6,12-dione) was synthesized (1.2 mg, 3.5 % yield) using 6-methoxy-1H-3,1-benzoxazine-2,4-dione as a starting material and the analogous procedure described for Compound **5**. ESI-MS m/z [M+H]+ calc'd for $C_{16}H_9ClN_2O_3$ 312.0; found 313.0.

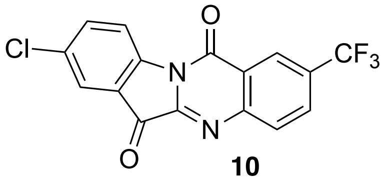

**10**

Compound **10** (8-chloro-2-(trifluoromethyl)indolo[2,1-b]quinazoline-6,12-dione) was synthesized (7.9 mg, 19 % yield) using 6-(trifluoromethyl)-1H-3,1-benzoxazine-2,4-dione as a starting material and the analogous procedure described for Compound **5**. ESI-MS m/z [M+H]+ calc'd for $C_{16}H_6ClF_3N_2O_2$ 350.0; found 350.9.

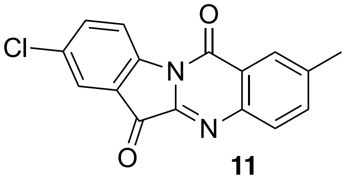

**11**

Compound **11** (8-chloro-2-methyl-indolo[2,1-b]quinazoline-6,12-dione) was synthesized (9.3 mg, 27 % yield) using 6-methyl-1H-3,1-benzoxazine-2,4-dione as a starting material and the analogous procedure described for Compound **5**. ESI-MS m/z [M+H]+ calc'd for $C_{16}H_9ClN_2O_2$ 296.0; found 297.00.

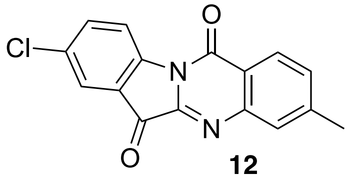

**12**

Compound **12** (8-chloro-3-methyl-indolo[2,1-b]quinazoline-6,12-dione) was synthesized (2.4 mg, 7.3 % yield) using 7-methyl-1H-3,1-benzoxazine-2,4-dione as a starting material and the analogous procedure described for Compound **5**. ESI-MS m/z [M+H]+ calc'd for $C_{16}H_9ClN_2O_2$ 296.0; found 297.0.

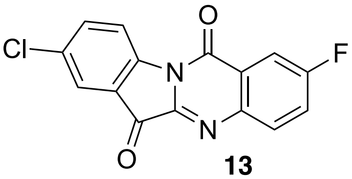

**13**

Compound **13** (8-chloro-2-fluoro-indolo[2,1-b]quinazoline-6,12-dione) was synthesized (3.9 mg, 11 % yield) using 6-fluoro-1H-3,1-benzoxazine-2,4-dione as a starting material and the analogous procedure described for Compound **5**. $^{1}H$

NMR (400 MHz, chloroform-*d*) δ ppm 7.36 - 7.42 (m, 1 H) 7.55 - 7.59 (m, 1 H) 7.68 - 7.72 (m, 1 H) 7.84 - 7.92 (m, 2H) 8.39 - 8.42 (m, 1 H). ESI-MS m/z [M+H]+ calc'd for $C_{15}H_6ClFN_2O_2$ 300.0; found 301.0.

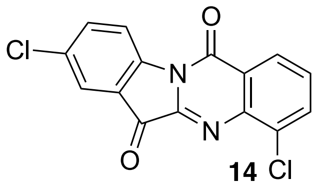

**14**

Compound **14** (4,8-dichloroindolo[2,1-b]quinazoline-6,12-dione) was synthesized (7.3 mg, 20 % yield) using 8-chloro-1H-3,1-benzoxazine-2,4-dione as a starting material and the analogous procedure described for Compound **5**. ESI-MS m/z [M+H]+ calc'd for $C_{15}H_6Cl_2N_2O_2$ 316.0; found 316.9.

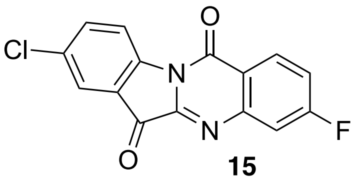

**15**

Compound **15** (8-chloro-3-fluoro-indolo[2,1-b]quinazoline-6,12-dione) was synthesized (12.6 mg, 36 % yield) using 7-fluoro-1H-3,1-benzoxazine-2,4-dione as a starting material and the analogous procedure described for Compound **5**. $^1$H NMR (400 MHz, chloroform-*d*) δ ppm. 7.18 - 7.25 (m, 1 H) 7.47 - 7.53 (m, 1 H) 7.55 - 7.59 (m, 1 H) 7.67 - 7.71 (m, 1 H) 8.24 - 8.30 (m, 1H) 8.37 - 8.42 (m, 1 H). ESI-MS m/z [M+H]+ calc'd for $C_{15}H_6ClFN_2O_2$ 300.0; found 301.0.

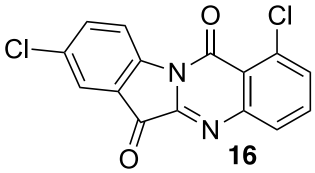

**16**

Compound **16** (1,8-dichloroindolo[2,1-b]quinazoline-6,12-dione) was synthesized (1.1 mg, 3.1 % yield) using 5-chloro-1H-3,1-benzoxazine-2,4-dione as a starting material and the analogous procedure described for Compound **5**. ESI-MS m/z [M+H]+ calc'd for $C_{15}H_6Cl_2N_2O_2$ 316.0; found 316.9.

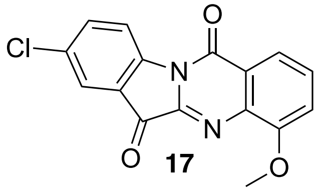

17

Compound **17** (8-chloro-4-methoxy-indolo[2,1-b]quinazoline-6,12-dione) was synthesized (3.9 mg, 11 % yield) using 8-methoxy-1H-3,1-benzoxazine-2,4-dione as a starting material and the analogous procedure described for Compound **5**. ESI-MS m/z [M+H]+ calc'd for $C_{16}H_9ClN_2O_3$ 312.0; found 313.0.

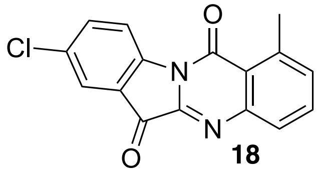

18

Compound **18** (8-chloro-1-methyl-indolo[2,1-b]quinazoline-6,12-dione) was synthesized using 5-methyl-1H-3,1-benzoxazine-2,4-dione as starting material using the analogous procedure described for Compound **5**. 1H NMR (400 MHz, chloroform-*d*) δ ppm 2.97 (s, 3 H) 7.44 (d, J=7.53 Hz, 1 H) 7.68 - 7.78 (m, 2 H) 7.86 - 7.90 (m, 2 H) 8.62 (d, J=8.53 Hz, 1 H). ESI-MS m/z [M+H]+ calc'd for $C_{16}H_9ClN_2O_2$ 296.04; found 296.90.

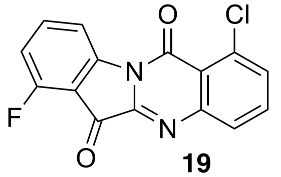

19

Compound **19** (1-chloro-7-fluoro-indolo[2,1-b]quinazoline-6,12-dione) was synthesized as follows: a solution of 5-chloro-1H-3,1-benzoxazine-2,4-dione (1.00 eq, 20 mg, 0.101 mmol), Rose Bengal (0.0500 eq, 5.2 mg, 0.00506 mmol) and DMA (0.9000 mL) was added to a 4 mL vial containing 4-fluoroindoline-2,3-dione (1.2 eq, 26.1 mg, 0.12 mmol) and potassium carbonate (1.00 eq, 14 mg, 0.101 mmol). The resulting mixture was stirred (uncapped) at ambient temperature under a 100 lumen white LED light (Milwaukee Tool, Brookfield, WI) for 18h. Purification by HPLC and SFC were performed as for Compound **5**. $^{1}$H NMR (400 MHz, methanol-$d_4$) δ ppm 7.12 (t, J=8.41 Hz, 1 H) 7.68 - 7.72 (m, 1 H) 7.72 - 7.78 (m, 1 H) 7.81 (td, J=8.31, 5.46 Hz, 1 H) 7.99 (dd, J=7.91, 1.38 Hz, 1 H) 8.52 (d, J=8.03 Hz, 1 H). ESI-MS m/z [M+H]+ calc'd for $C_{15}H_6ClFN_2O_2$ 300.01; found 300.90.

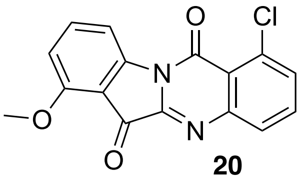

**20**

Compound **20** (1-chloro-7-methoxy-indolo[2,1-b]quinazoline-6,12-dione) was synthesized using 4-methoxyindoline-2,3-dione as starting material using the analogous procedure described for Compound **19**. 1H NMR (400 MHz, chloroform-*d*) δ ppm 4.10 (s, 3 H) 6.94 (d, *J*=8.53 Hz, 1 H) 7.64 - 7.77 (m, 3 H) 7.98 (dd, *J*=7.97, 1.32 Hz, 1 H) 8.28 (d, *J*=7.91 Hz, 1 H). ESI-MS m/z [M+H]+ calc'd for $C_{16}H_9ClN_2O_3$ 312.03; found 313.00.

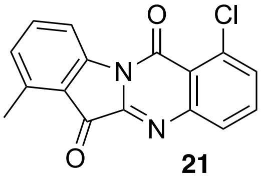

**21**

Compound **21** (1-chloro-7-methyl-indolo[2,1-b]quinazoline-6,12-dione) was synthesized using 4-methyl-2,3-dihydro-1H-indole-2,3-dione as starting material using the analogous procedure described for Compound **19**. $^{1}$H NMR (400 MHz, methanol-$d_4$) δ ppm 2.77 (s, 3 H) 7.22 (d, J=7.78 Hz, 1 H) 7.62 - 7.70 (m, 2 H) 7.70 - 7.75 (m, 1 H) 7.97 (dd, J=7.97, 1.32 Hz, 1 H) 8.53 (d, J=8.03 Hz, 1 H). ESI-MS m/z [M+H]+ calc'd for $C_{16}H_9ClN_2O_2$ 296.04; found 297.00.

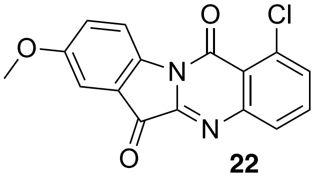

**22**

Compound **22** (1-chloro-8-methoxy-indolo[2,1-b]quinazoline-6,12-dione) was synthesized using 5-methoxy-2,3-dihydro-1H-indole-2,3-dione as starting material using the analogous procedure described for Compound **19**. 1H NMR (400 MHz, chloroform-*d*) δ ppm 3.92 (s, 3 H) 7.33 (d, *J*=8.28 Hz, 1 H) 7.40 (d, *J*=2.76 Hz, 1 H) 7.65 - 7.69 (m, 1 H) 7.69 - 7.75 (m, 1 H) 7.96 (d, *J*=7.75 Hz, 1 H) 8.58 (d, *J*=8.78 Hz, 1 H). ESI-MS m/z [M+H]+ calc'd for $C_{16}H_9ClN_2O_3$ 312.03; found 312.95.

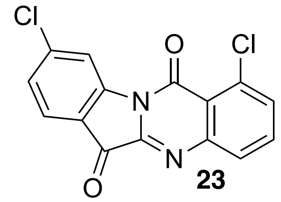

**23**

Compound **23** (1,9-dichloroindolo[2,1-b]quinazoline-6,12-dione) was synthesized using 6-chloroindoline-2,3-dione as starting material using the analogous procedure described for Compound **19**. $^{1}$H NMR (400 MHz, chloroform-*d*) δ ppm 7.44 (dd, *J*=8.16, 1.76 Hz, 1 H) 7.66 (dd, *J*=8.53, 2.01 Hz, 1 H) 7.87 (d, *J*=8.03 Hz, 1 H) 8.02 (d, *J*=2.01 Hz, 1 H) 8.38 (d, *J*=8.53 Hz, 1 H) 8.68 (d, *J*=1.76 Hz, 1 H). ESI-MS m/z [M+H]+ calc'd for $C_{15}H_{6}Cl_{2}N_{2}O_{2}$ 315.98; found 316.95.

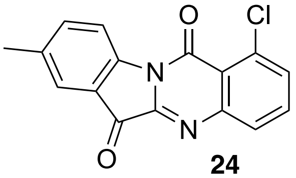

**24**

Compound **24** (1-chloro-8-methyl-indolo[2,1-b]quinazoline-6,12-dione) was synthesized using 5-methyl-2,3-dihydro-1H-indole-2,3-dione as starting material using the analogous procedure described for Compound **19**. $^{1}$H NMR (400 MHz, chloroform-d) δ ppm 2.48 (s, 3 H) 7.62 (d, J=8.16 Hz, 1 H) 7.65 - 7.69 (m, 1 H) 7.69 - 7.75 (m, 2 H) 7.97 (dd, J=7.91, 1.38 Hz, 1 H) 8.55 (d, J=8.28 Hz, 1 H). ESI-MS m/z [M+H]+ calc'd for $C_{16}H_{9}ClN_{2}O_{2}$ 296.04; found 297.00.

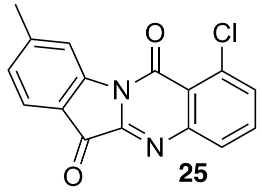

**25**

Compound **25** 1-chloro-9-methyl-indolo[2,1-b]quinazoline-6,12-dione was synthesized using 6-methylindoline-2,3-dione as starting material using the analogous procedure described for Compound **19**. $^{1}$H NMR (400 MHz, methanol-$d_4$) δ ppm 2.56 (s, 3 H) 7.19 - 7.26 (m, 1 H) 7.60 - 7.68 (m, 1 H) 7.68 - 7.75 (m, 1 H) 7.81 (d, J=7.78 Hz, 1 H) 7.96 (d, J=7.98 Hz, 1 H) 8.50 - 8.55 (m, 1 H). ESI-MS m/z [M+H]+ calc'd for C16H9ClN2O2 296.04; found 296.95.

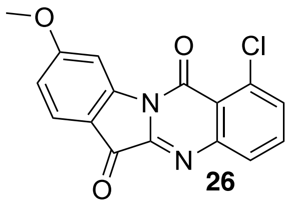
**26**

Compound **26** (1-chloro-9-methoxy-indolo[2,1-b]quinazoline-6,12-dione) was synthesized using 6-methoxyindoline-2,3-dione as starting material using the analogous procedure described for Compound **19**. [1]H NMR (400 MHz, chloroform-*d*) δ ppm 4.04 (s, 3 H) 6.92 (dd, *J*=8.53, 2.26 Hz, 1 H) 7.67 (dd, *J*=7.97, 1.32 Hz, 1 H) 7.73 (t, *J*=7.97 Hz, 1 H) 7.87 (d, *J*=8.53 Hz, 1 H) 7.97 (dd, *J*=7.91, 1.25 Hz, 1 H) 8.25 (d, *J*=2.13 Hz, 1 H). ESI-MS m/z [M+H]+ calc'd for $C_{16}H_9ClN_2O_3$ 312.03; found 312.95.

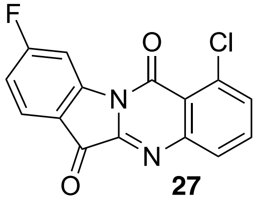
**27**

Compound **27** (1-chloro-9-fluoro-indolo[2,1-b]quinazoline-6,12-dione) was synthesized using 6-fluoro-2,3-dihydro-1H-indole-2,3-dione as starting material using the analogous procedure described for Compound **19**. [1]H NMR (400 MHz, chloroform-*d*) δ ppm 7.15 (td, J=8.47, 2.26 Hz, 1 H) 7.68 - 7.72 (m, 1 H) 7.76 (t, J=7.60 Hz, 1 H) 7.95 - 8.00 (m, 2 H) 8.45 (d, J=2.13 Hz, 1 H). ESI-MS m/z [M+H]+ calc'd for $C_{15}H_6ClFN_2O_2$ 300.01; found 301.00.

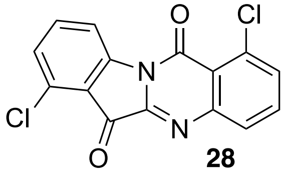
**28**

Compound **28** (1,7-dichloroindolo[2,1-b]quinazoline-6,12-dione) was synthesized using 4-chloroindoline-2,3-dione as starting material using the analogous procedure described for Compound **19**. [1]H NMR (400 MHz, chloroform-*d*) δ ppm 7.41 (d, J=8.16 Hz, 1 H) 7.68 - 7.78 (m, 3 H) 7.99 (dd, J=7.91, 1.25 Hz, 1 H) 8.66 (d, J=8.03 Hz, 1 H). ESI-MS m/z [M+H]+ calc'd for $C_{15}H_6Cl_2N_2O_2$ 315.98; found 316.90.

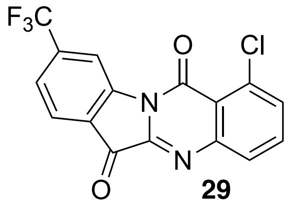

**29**

Compound **29** (1-chloro-9-(trifluoromethyl)indolo[2,1-b]quinazoline-6,12-dione) was synthesized using 6-(trifluoromethyl)indoline-2,3-dione as starting material using the analogous procedure described for Compound **19**. ESI-MS $[M+H]^+$ calc'd for $C_{16}H_6ClF_3N_2O_2$, 350.0; found, 350.90.

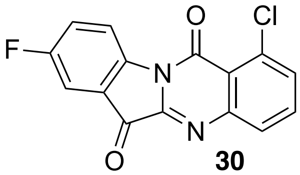

**30**

Compound **30** (1-chloro-8-fluoro-indolo[2,1-b]quinazoline-6,12-dione) was synthesized using 5-fluorisatin as starting material using the analogous procedure described for Compound **19**. $^1H$ NMR (400 MHz, chloroform-*d*) δ ppm 7.51 (td, J=8.66, 2.76 Hz, 1 H) 7.61 (dd, J=6.53, 2.76 Hz, 1 H) 7.68 - 7.72 (m, 1 H) 7.72 - 7.77 (m, 1 H) 7.98 (dd, J=7.91, 1.38 Hz, 1 H) 8.71 (dd, J=8.85, 4.08 Hz, 1 H) . ESI-MS m/z [M+H]+ calc'd for $C_{15}H_6ClFN_2O_2$ 300.01; found 300.95.

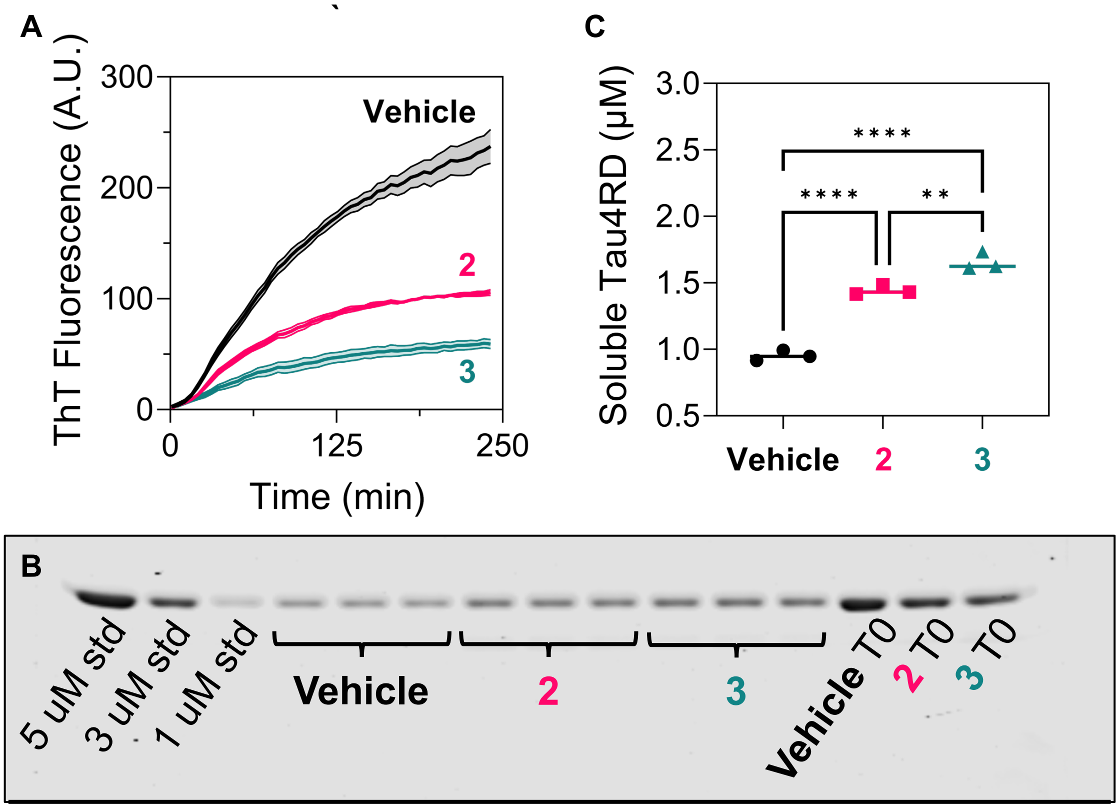

Figure S1: Confirmation of TA analog activity by label-free centrifugation assays. A) Tau4RD (5 µM) aggregation kinetics in the presence of 5 µM 3 kDa heparin and either 10 µM **2**, 5 µM **3**, or vehicle control, in filtered buffer containing 20 mM Tris, 50 mM NaCl, and 1 mM TCEP. ($n$ = 3 or 4, error bands show mean +/- SEM.) B) SDS-PAGE gel densitometry of soluble tau4RD remaining at $t$ = 240 min from panel A, including additional Tau4RD concentration standards. C) Soluble tau4RD remaining at $t$ = 240 min calculated from the results of panel B.

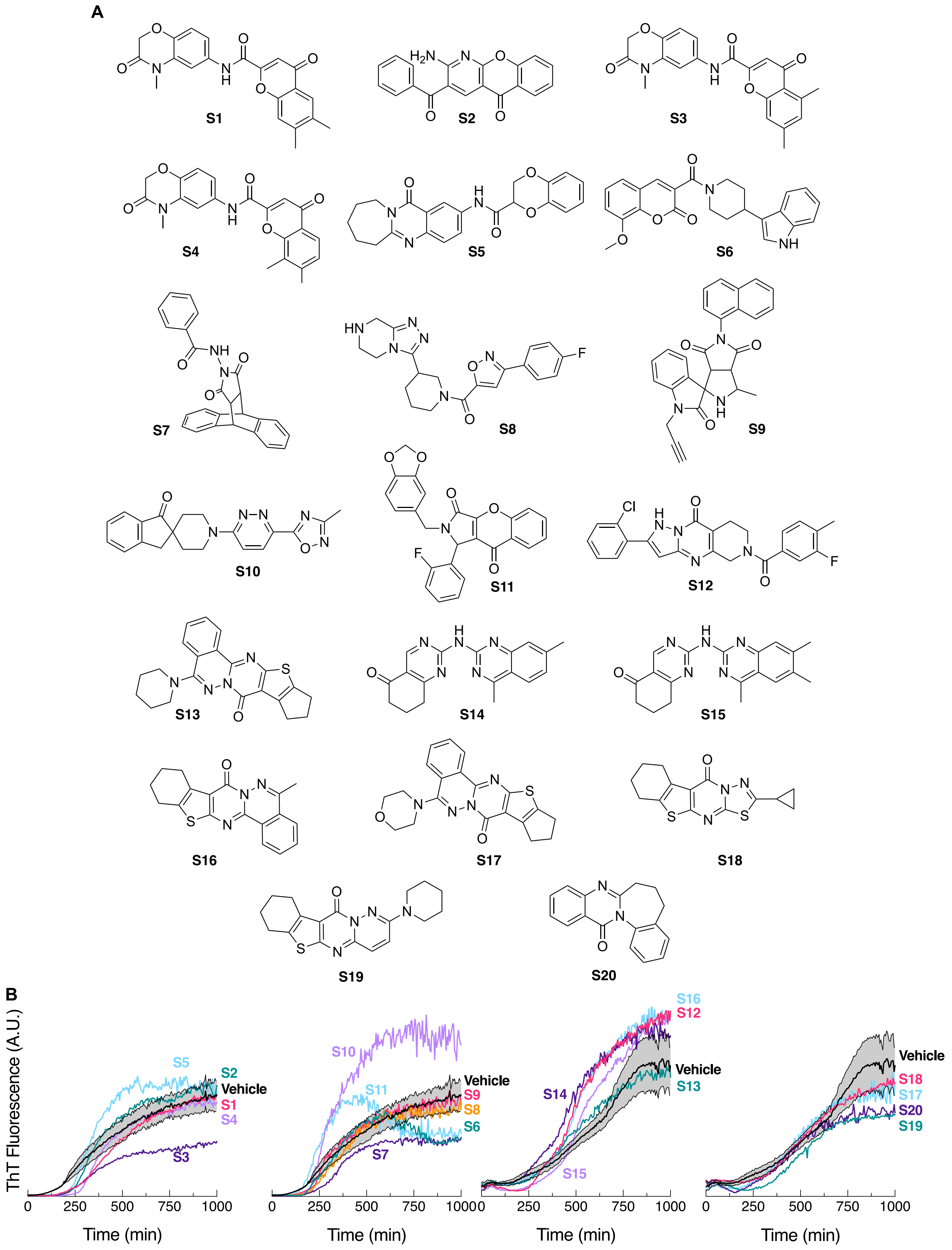

Figure S2: Structures (A) and aggregation inhibition activity (B) of non-tryptanthrin compounds in screen. Conditions and formatting match those for Fig. 1.

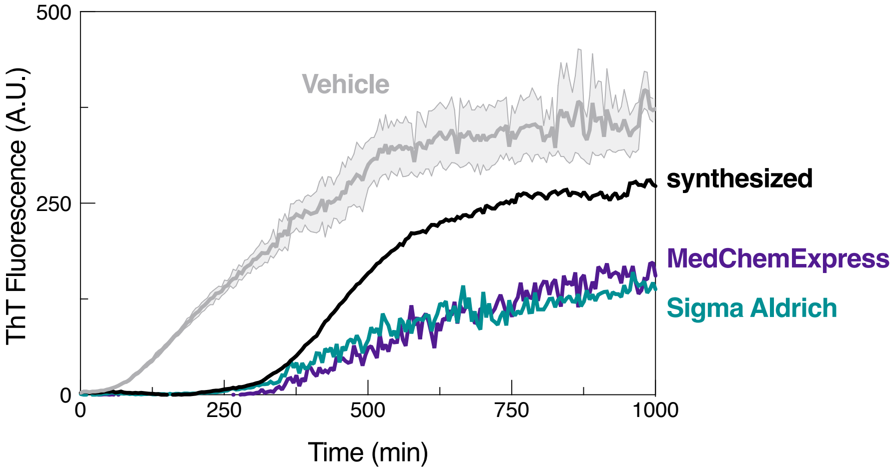

Figure S3: Comparison of synthesized vs. commercially available tryptanthrin (**1**). Aggregation of 5 μM tau4RD is inhibited to similar extents by 0.31 μM **1** synthesized in-house or acquired from Sigma Aldrich (St. Louis, MO) or MedChemExpress (Monmouth Junction, NJ).

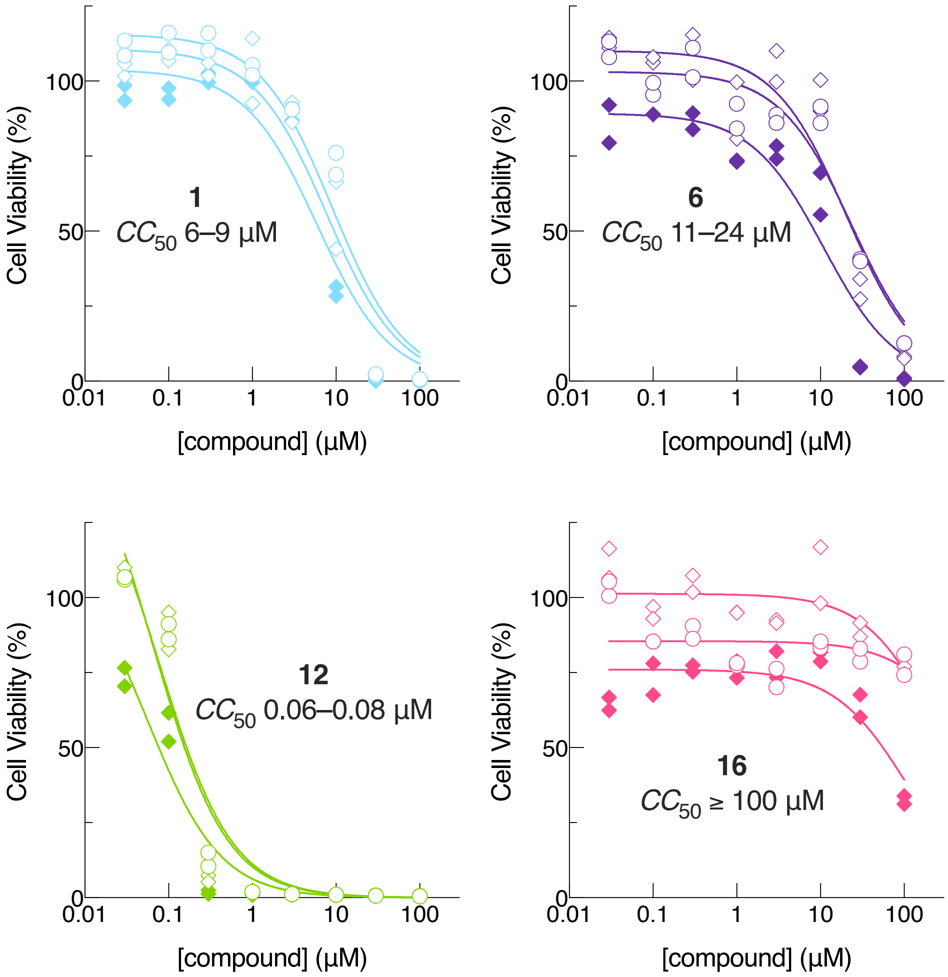

Figure S4: Cytotoxicity of selected TA analogs towards HepG2 human hepatocellular carcinoma cells in culture. Cytotoxicity assays were conducted by Eurofins Discovery Services (Fremont, CA) using the following protocol: cells were plated in 384-well plates at a density of 3000 cells/well, cultured overnight at 37ºC, and transferred into assay media. These media included TA analogs at concentrations ranging from 0.03–100 µM in modified Eagle's medium without phenol red, 1% fetal bovine serum, 2 mM L-glutamine, 50 U/mL penicillin, 50 µg/mL streptomycin, supplemented with glucose (circles) or galactose (diamonds). Cell viability was measured in duplicate after 24 h (open symbols) or 72 h (closed symbols) using the Cell Titer Glo assay (Promega Corp., Madison, WI).
